# *In vitro* dormancy models improve ability to predict treatment response in severe marmoset tuberculosis lesions

**DOI:** 10.64898/2026.05.13.724664

**Authors:** Joshua J. Whiteley, Talia Greenstein, Mariana Pereira Moraes, Maral Budak, Danielle M. Weiner, Ayan Abdi, Joel D. Fleegle, Felipe Gomez, Laura E. Via, Clifton E. Barry, Jansy Sarathy, Denise Kirschner, Veronique Dartois, Bree B. Aldridge

**Author notes:** These authors contributed equally to this work (Joshua J. Whiteley, Talia Greenstein, Mariana Pereira Moraes).

## Abstract

Tuberculosis lesions are structurally and physiologically heterogeneous, creating microenvironments that restrict antibiotic penetration and alter bacterial susceptibility. This heterogeneity remains a major impediment to shortening tuberculosis treatment, as it is difficult to predict which regimens will sterilize bacteria in these hard-to-treat lesion niches. Many *in vitro* systems and commonly used animal models fail to recapitulate the necrotic, caseous lesion pathology associated with treatment failure and relapse. Here, we show that resource-efficient *in vitro* assays engineered to mimic key lesion microenvironments, including those with lipid-rich, caseum-like matrices with controlled oxygenation and pH, generate drug-response metrics that predict lesion-level treatment responses in a non-human primate model. We stratified individual marmoset lesions by baseline 2-deoxy-2-[18F] fluoro-D-glucose ([18F] FDG) positron emission tomography/computed tomography (PET/CT) features, then quantified drug potency and interactions across three lipid-induced dormancy (LIDs) conditions under both equipotent and lesion pharmacokinetic-informed dosing schemes. LIDs-based and pharmacokinetic-informed measurements aligned more closely with treatment responses in severe lesions than conventional *in vitro* data and recapitulated interaction patterns consistent with known regimen performance. Integrating imaging-derived and *in vitro* features in multivariate, sequential computational models improved predictive accuracy over baseline imaging alone and identified a subset of informative *in vitro* metrics. Together, these results establish a scalable framework linking lesion-mimicking assays to lesion-specific outcomes, enabling earlier, more cost-effective prioritization of pre-clinical regimens aimed at sterilizing hard-to-treat tuberculosis lesions and shortening therapy.

## Introduction

The hallmark of pulmonary tuberculosis (TB) is the formation of diverse lesion types, including cellular granulomas, necrotic caseous granulomas, and cavities. Lesions vary in size, architecture, and microenvironmental composition (i.e., lipid content, metal availability, oxygen tension, and pH) (Sawyer, 2023; Kurthkoti, 2017; Via, 2008; Baker, 2014). Severe TB is characterized by the presence of extensive necrotic, caseous, and cavitary lesions, where limited drug penetration (a pharmacokinetic feature) and reduced bacterial susceptibility (a pharmacodynamic feature) in resident bacterial populations contribute to the need for prolonged patient therapy and increased relapse risk (Dartois, 2025; Prideaux, 2015; Ordonez, 2020; Imperial, 2018; Kim, 2021). As *Mycobacterium tuberculosis* (Mtb) adapts to distinct lesion environments, the bacteria shift metabolic state and growth rate, giving rise to lesion-dependent drug susceptibilities (Barry, 2009; Harper, 2012; Sarathy, 2018). Reaching and sterilizing each of these bacterial subpopulations requires a diverse combination of drugs to penetrate the different lesion structures and to target the physiologically distinct bacterial populations within them (Greenstein, 2023; Cicchese, 2020; Kerantzas, 2017; Waddell, 2015). Necrotic, caseous, and cavitary granulomas are a critical focal point when designing new regimens with the intent to shorten TB therapy.

The six-month duration of standard drug therapy is to ensure cure in patients who have extensive necrotic, caseous, or cavitary disease (Barry, 2009; Imperial, 2018). For patients with severe pathology, prolonged therapy remains necessary to achieve cure without relapse. Because the six-month regimen is the clinical standard treatment, patients with less severe disease are consequently overtreated (Imperial, 2018), and patients may find treatment adherence difficult due to several factors, including side effects. Developing shorter treatments, then, is a vital step towards ensuring proper treatment for more patients. A key to shortening treatment lies in designing regimens that effectively target Mtb within hard-to-treat lesions (Dartois, 2025; Prideaux, 2015; Sarathy, 2018). Designing these regimens requires experimental models that recapitulate the diverse microenvironments of severe granulomas, enabling evaluation of drug penetration and efficacy under clinically relevant conditions (Sarathy, 2018; Rifat, 2018; Sarathy, 2023). Preclinical systems that preserve lesion-level pathology and treatment response are therefore essential for evaluating regimens designed to shorten therapy in severe disease.

Non-human primate (NHP) models are thought to most closely resemble the complex lesion pathology and severity of TB disease in humans (Via, 2013; Lin et al., 2013; Mehra et al., 2010; Sharan et al., 2022). NHP models are also compatible with 2-deoxy-2-[18F]fluoro-D-glucose (FDG) positron emission tomography/computed tomography (PET/CT) imaging, which enables longitudinal, lesion-level quantification of disease progression and treatment response. Using PET/CT imaging, disease progression and resolution can be tracked over time. Multivariate profiles from per-lesion PET/CT analyses in drug-treated Mtb-infected marmosets have been shown to align with clinical outcomes (Greenstein, 2026). Unfortunately, NHP imaging studies are costly and space-intensive, limiting their scalability and making them impractical for large-scale or high-throughput applications despite their strong translational relevance. These constraints underscore the need for more accessible, complementary systems capable of capturing lesion-specific treatment dynamics.

Together, these limitations motivate complementary computational approaches that can leverage limited, information-rich datasets to predict lesion-specific treatment dynamics and guide regimen design. Predictive modeling has become an increasingly powerful approach for translating preclinical data into regimen design and clinical outcomes. Multiple groups have developed computational frameworks that link pharmacokinetics, bacterial dynamics, and lesion heterogeneity to treatment response, including pharmacokinetic modeling work from the Savic laboratory (Goh, 2025; Strydom, 2019), systems-level modeling from the Kirschner and Chandrasekaran laboratories that captures granuloma-scale and metabolic interactions (Cicchese, 2021), and PK/PD modeling for regimen design from Gumbo and Hermann (Gumbo, 2022). We have developed computational models that correlate *in vitro* drug-response metrics to relapse outcomes in the BALB/c relapsing mouse model (Larkins-Ford, 2021; Larkins-Ford, 2022); however, those systems lack the necrotic and caseous pathology that defines severe disease.

Here, we address this gap by integrating lesion-focused imaging with resource-sparing *in vitro* assays to predict outcomes at the lesion level. We first classify individual marmoset lesions according to baseline PET/CT features, then measure drug responses in lipid-induced dormancy (LIDs) assays that simulate key attributes of hard-to-treat environments (such as lipid-rich caseum, oxygen, and pH gradients) under equipotent and lesion PK-informed dosing. By correlating these readouts with per-lesion PET/CT trajectories, we identify a set of *in vitro* metrics that better recapitulate severe lesion responses than conventional *in vitro* assays. Finally, we combine imaging and *in vitro* data in multivariate sequential models to predict lesion-level progression. Together, these methods establish a scalable framework that links lesion-mimicking assays to lesion-specific outcomes, enabling earlier and lower-cost prioritization of regimens aimed at sterilizing hard-to-treat lesions and shortening TB therapy.

## Results

### Marmoset lesions can be categorized by severity using quantitative PET/CT data

In aiming to develop resource-sparing *in vitro* models to model lesion-specific treatment outcomes, we leveraged PET/CT data from the marmoset model that provided lesion-level resolution and trajectory during treatment (Greenstein, 2026). The first step was to categorize lesion-specific datasets by severity type. We previously observed that marmoset lesions with severe pathologies, as determined by PET/CT, were more likely to be cavitary or necrotic than those with low or moderate PET/CT pathologies (Greenstein, 2026). Severity designations were based on PET inflammatory activity and CT lesion size and density (Greenstein, 2026). We hypothesized that baseline (start-of-treatment) PET/CT features, including the mean FDG standard uptake value, mean radiodensity, and tissue volumes, could be used to classify lesions as more or less severe.

To test this hypothesis, we analyzed lesion-specific treatment outcomes in a marmoset model. Our dataset consisted of marmosets infected with *Mycobacterium tuberculosis* and assigned to one of 22 treatment arms, with five marmosets per arm (Budak, 2024; Greenstein, 2026). Of these treatment arms, nine were single drug regimens; the remaining 13 were combinations of at least two and up to four antibiotics (Table 1). Treatment began 6 to 7 weeks after infection. Treatment response was monitored by PET/CT at treatment start and after 2, 4, 6, and 8 weeks of treatment, with week 8 corresponding to end-of-treatment necropsy and CFU enumeration.

**Table 1.** Abbreviations used in this study and brief descriptions of said abbreviations.

| Abbreviation | meaning |
| --- | --- |
| *antibiotics* |  |
| B | bedaquiline |
| D | delamanid |
| H | isoniazid |
| L | linezolid |
| M | moxifloxacin |
| Q | quabodepistat |
| Pa | pretomanid |
| R | rifampicin |
| Z | pyrazinamide |
| *in vitro metrics* |  |
| FxC<sub>n</sub> | log2 scaled inhibitory **or** bactericidal concentration at n % |
| E<sub>inf</sub> | the effect at infinite drug concentration |
| GR<sub>max</sub> | normalized growth inhibition effect at maximum dose |
| t<sub>c</sub> | constant time point |
| t<sub>T</sub> | terminal time point |

These longitudinal scans provided lesion-level measurements of FDG uptake as a readout of inflammatory activity, CT radiodensity as a readout of lesion density and composition, and soft and hard tissue volumes as measures of lesion burden. Full imaging, ROI, and necropsy workflows are described in Methods and Greenstein (2026).

We set out to categorize these lesions by severity, as determined by the PET/CT measurements at the start of treatment. Lesion clustering was performed by unsupervised clustering using Leiden community detection, resulting in 16 clusters (Figure 1A and Methods). Standard deviation of radiodensity was calculated as a proxy for lesion composition; other calculated feature values were the median values for mean PET activity, mean radiodensity, hard volume, and soft volume (Table S1). We then performed hierarchical clustering on the medians to identify relationships (Figure 1B and Methods). Hierarchical clustering revealed two distinct groups in the top clade, which evenly split the dataset into two groups of eight clusters each (Figure 1B). These two groups represented a more severe (n=566 lesions) and a less severe (n=627 lesions) set of lesions (Figure 1C).

**Figure 1.**
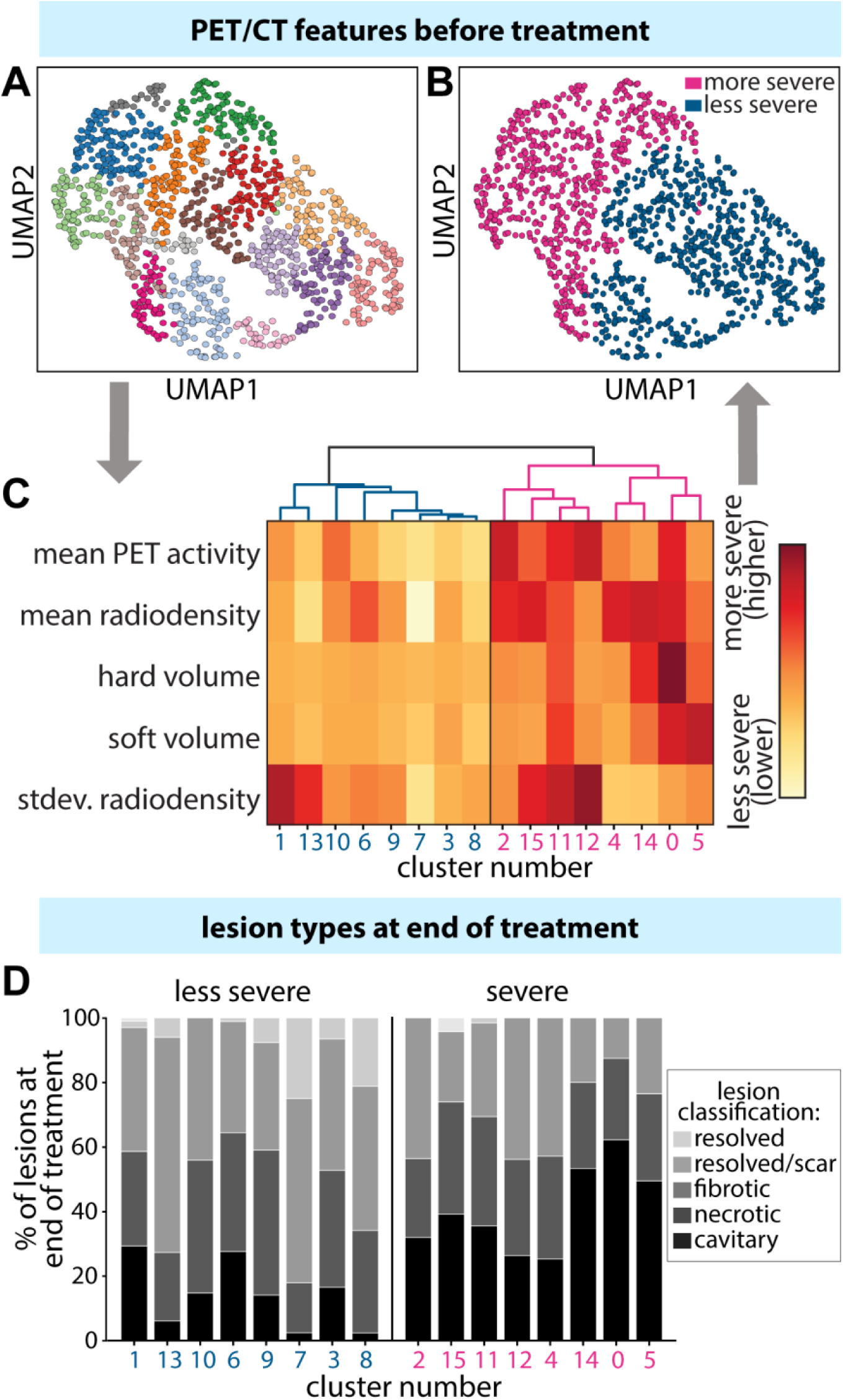
Marmoset TB lesions can be categorized into severe and less severe classes based on PET/CT measurements prior to treatment start. (A) UMAP of the results of Leiden clustering on individual lesions (n = 1193), based on mean PET activity, mean radiodensity, hard volume, soft volume, and standard deviation of radiodensity. Each color represents a unique cluster (n=16). Leiden was run on a kNN graph built from PCA features (cosine distance). (B) Hierarchical clustering and heatmap of the median feature value across communities from the Leiden community detection. Blue (left branch) indicates the less severe clusters, and red (right branch) indicates the more severe clusters. Hierarchical clustering used cosine distance and average linkage. (C) Severe and less severe labels applied to the UMAP of individual lesions from A, determined by the clustering in B. (D) Distributions of lesion type at the end of eight weeks of treatment across less and more severe lesions at start. Severe lesions have higher proportions of necrotic and cavitary lesions (severe lesion types) at the end of treatment, whereas less severe lesions have higher proportions of fibrotic (less severe) and resolved lesions.

To confirm the lesion severity classifications assigned based on the start-of-treatment PET/CT data, we compared them with histopathology evaluations of each lesion at the end of treatment. Lesions were classified by end-of-treatment severity as less severe (fibrotic), severe (necrotic), severe (cavitary), and resolved (absent or scar only). Each cluster group contained the full spectrum of lesion types, although their relative abundances varied within each subgroup. There was a higher abundance of less severe and fully resolved lesions in the less severe cluster group, and a higher proportion of severe lesion types (both necrotic and cavitary) in the severe cluster group (Figure 1D). The groupings support our initial hypothesis that PET/CT features observed prior to treatment can be used to distinguish lesion severity and predict distinct treatment responses.

### Design of resource-sparing in vitro dose-response measurements to model lesion pharmacokinetics and pharmacodynamics

We previously developed a suite of *in vitro* models to predict relapse outcome after combination drug treatment in the BALB/c mouse, which develops cellular lesions, a less severe form of pathology (Larkins-Ford et al., 2021; Larkins-Ford et al., 2022). Modeling based on the BALB/c mouse data suggested that *in vitro* combination potency metrics (e.g., drug efficacy measures) predict *in vivo* treatment success. Given that these *in vitro* measurements used equipotency in the dosing strategy, these data may not accurately reflect how the pharmacokinetics (PK) of each drug in a combination determine the combination’s potency. In addition to including more host-relevant dosing ratios, we sought to include experimental conditions that better reflect the spectrum of microenvironments in severe lesion types. Insights from *ex vivo* caseum studies informed our design (Sarathy, 2018; Sarathy, 2023). *Ex vivo* caseum models highlighted the need for a dormancy-like, lipid-rich growth condition with varying oxygen tension and pH. We hypothesized that combining our pharmacokinetic dosing strategy (PK dosing) with more complex growth conditions would better reflect treatment responses in severe pathologies and, by extension, improve correlations between *in vitro* readouts and clinical outcomes.

To model the dormant, non-replicating state of Mtb in caseous lesions, we established dormancy conditions designed to reproduce the lipid-rich microenvironment characteristic of these lesions (Deb et al., 2009; Sarathy, 2020; Wayne & Sohaskey, 2001; Baker, Johnson, & Abramovitch, 2014; Cho et al., 2007; Piddington et al., 2000; Daniel et al., 2011). We developed three lipid-induced dormancy models (LIDs), each composed of a lipid-rich base that included cholesterol, arachidonic acid, linoleic acid, palmitic acid, oleic acid, and stearic acid. The lipid composition was guided by *ex vivo* caseum measurements, and the choice of neutral versus acidic pH and hypoxia reflects reported caseum physiology (Sarathy, 2023). The three models ultimately developed were combinations of pH (neutral or acidic) and oxygenation (hypoxic or normoxic): neutral pH, normoxic (7N); neutral pH, hypoxic (7H); and acidic pH, normoxic (5N). Mtb was acclimated to these conditions for four weeks to achieve a metabolically inactive, non-replicative viable state (Figure S1). After four weeks of acclimation, the bacteria were treated with antibiotics for two weeks. Following the two-week drug treatment, bacteria were transferred to charcoal agar plates to allow for a treatment-free recovery period. During recovery, we used a plate reader to measure the metabolic activity of live bacteria via luminescence facilitated by an autoluminescent reporter strain of Mtb (Gold et al., 2015; Larkins-Ford, 2021; Methods). These data were then organized into dose-response curves.

To benchmark the LIDs conditions against established lesion-relevant systems, we compared them with *ex vivo* caseum and an *in vitro* caseum surrogate. *Ex vivo* caseum assays use authentic caseous material from TB lesions and have shown that Mtb in lipid-rich, non-replicating environments exhibit marked drug tolerance and a distinct hierarchy of antibiotic susceptibility (Sarathy, 2018; Sarathy, 2020). The *in vitro* caseum surrogate was developed to reproduce key features of caseum-like lipid matrices in a more scalable experimental format (Sarathy, 2023). We therefore used these assays as a reference system to assay whether the LIDs conditions recapitulate drug-tolerance patterns observed in caseous lesion environments.

To evaluate the extent of drug tolerance induced in LIDs conditions, we measured the dose-response curves for 14 different anti-mycobacterial drugs. We determined the minimal bactericidal concentration required to kill 90% of a specific bacterial population (MBC90) for 14 drugs across the three LIDs conditions. We then compared these MBC90 values to (i) inhibitory concentrations required to inhibit 90% of growth (IC90) from non-LIDs growth assays and (ii) MBC90 values from other models. These comparison models included assays validated as predictive of outcomes in the RMM, such as the nitrate-induced dormancy assay, acidic growth medium, and conditions using cholesterol or butyrate as the sole carbon source (Larkins-Ford, 2021. Of the three LIDs conditions, hypoxic conditions resulted in greater tolerance than the two normoxic conditions. Across the assays tested, several conditions could be compared with the rabbit *ex vivo* caseum (Figure 2A, Table S2, Table S3). Among the three LIDs conditions, the neutral-normoxic assay most closely matched rabbit *ex vivo* caseum by cross-drug rank order and absolute potency, resulting in a ρ of 0.75, followed by the acidic-normoxic condition with a ρ of 0.65, and finally the neutral-hypoxic condition with a ρ of 0.57. Among the non-LIDs assays, nitrate-induced dormancy correlated more with *ex vivo* caseum and the caseum surrogate than with the conditions with an acidic medium, or those with butyrate or cholesterol as a sole carbon source. We base our conclusion on both the cross-drug susceptibility rank-ordering and the absolute potency levels observed in LIDs compared with *ex vivo* caseum. For example, cell-wall synthesis inhibitors exhibit low IC90s in simple assays but require substantially higher concentrations to reach MBC90 in the LIDs assays and in *ex vivo* caseum and caseum-surrogate assays (Figure 2A, top left vs. top right), consistent with the higher ρ values reported above. Similarities in drug tolerance across multiple antibiotics with different mechanisms of action suggest that one or more of the LIDs conditions could be a more suitable model of the dormant, drug-tolerant bacterial states that occur *in vivo*.

**Figure 2.**
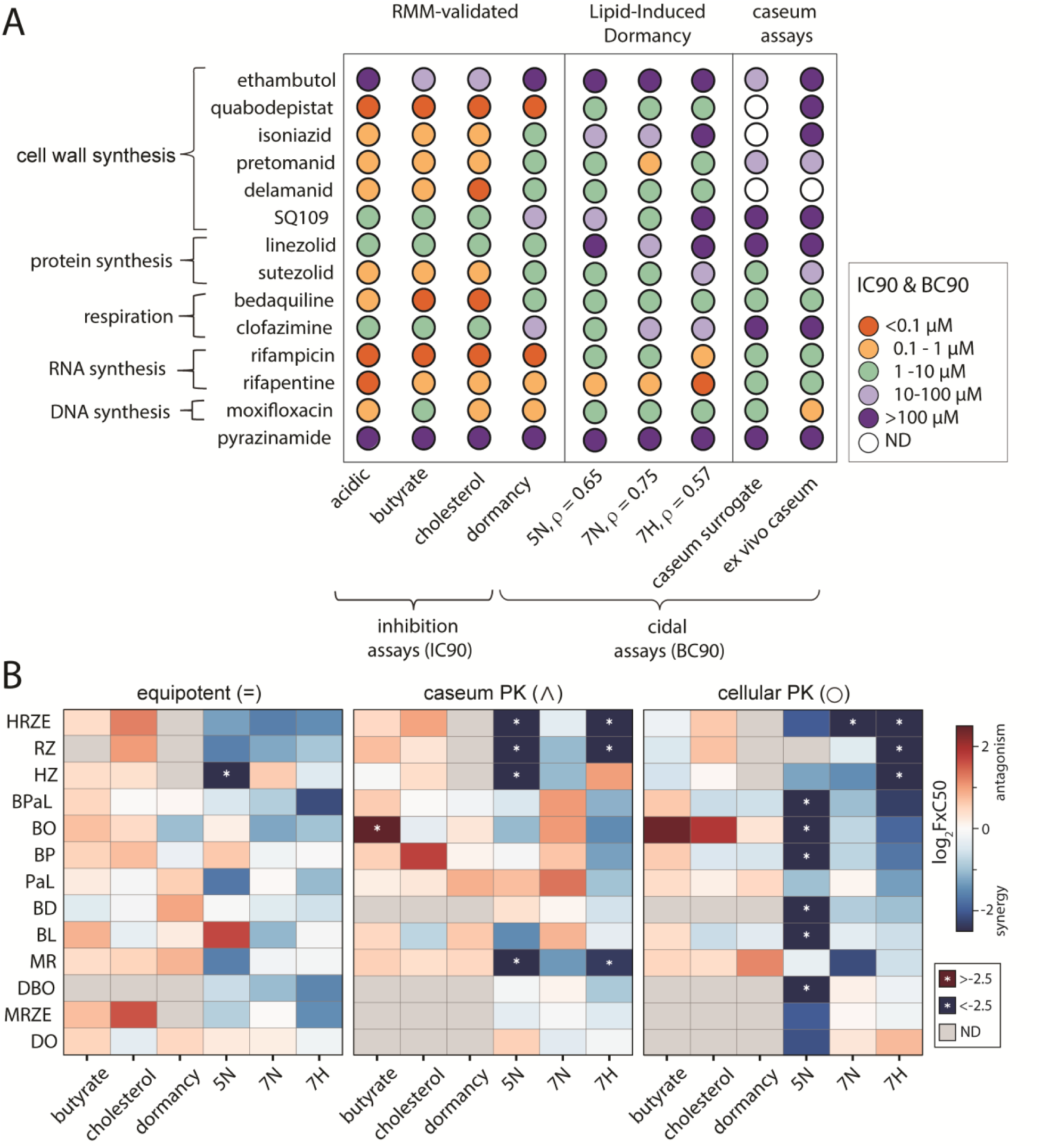
LIDs drug and drug combination responses. (A) single drug susceptibilities in RMM-validated experimental models (acidic, butyrate, cholesterol, dormancy), LIDs models (5N, 7N, 7H), and more advanced caseum models (caseum surrogate, *ex vivo* caseum). IC90 and BC90 values are reported for 14 anti-mycobacterial drugs with diverse underlying mechanisms. Rho values under the LIDs conditions are Spearman correlations and are calculated with respect to the *ex vivo* caseum values. (B) Heatmap comparing log2 FxC50 (FIC for butyrate, cholesterol, and simple dormancy; FBC for 5N, 7N, 7H) values across experimental conditions, drug combinations, and dosing strategies, sorted from lowest 7H FxC50 to highest. A white star on a heatmap tile indicates an FxC with a magnitude of greater than 2.5.

To determine whether drug interactions measured in LIDs models are redundant with more resource-sparing models, we compared the drug interaction profiles in the LIDs models with those from the RMM-validated *in vitro* models. We measured the 13 drug combinations for which we have lesion-specific treatment information in the marmoset. Using equipotent dosing (Figure 2B, left), we observed poor correlation between the FIC50s measured RMM-validated *in vitro* and the FBC50s measured in LIDs conditions. We also observed that drug interactions in LIDs models were far more synergistic across the suite of drug combinations tested than in the RMM-validated models. Taken together, we conclude that the LIDs conditions provide unique information that is not obtainable from simple *in vitro* assays.

In the equipotent dosing strategy, drugs were combined so that each drug contributed an equivalent potency level (either IC90 or MBC90), aligned to the seventh of ten doses in the dose-response series. Recognizing that drugs penetrate lesions at different concentrations based on lesion composition and structure, we also tested drug combination responses at dosing ratios reflecting caseum and cellular lesion pharmacokinetics derived from simulations from the GranSim model (Box S1, Table S5). To design PK-informed DiaMOND experiments, we implemented a dosing strategy centered around the simulated C_max_, the peak concentration of a drug in a compartment, on a per-drug basis (Methods). All three dosing strategies were measured across all LIDs models and the three chosen non-LIDs conditions. No LIDs condition showed a Spearman correlation with a p-value of >0.05 between dosing strategies. The range of intra-condition ρ values was −0.31 to 0.52, and the range of p-values was between 0.02 and 0.65 (Table S4). We conclude that none of the tested LIDs conditions are redundant.

Across LIDs conditions, PK-informed dosing revealed more synergistic relationships than equipotent dosing (Figure 2B), especially in the neutral-hypoxic (7H) condition. PaL, for instance, shifted from antagonism to additivity to synergy depending on condition and dosing strategy, highlighting that each LIDs assay captures distinct interaction behavior. At the regimen level, HRZE exhibited the most consistently positive interactions across conditions, whereas MRZE showed much weaker interaction effects. This mirrors clinical evidence that MRZE is inferior to HRZE in humans (Dorman, 2009; Jawahar, 2013; Gillespie, 2014). Together, these results indicate that PK-dosed LIDs assays provide nonredundant information relative to the RMM-validated models.

### A subset of in vitro metrics from multiple models correlates with PET/CT outcomes in marmosets

We next asked whether these *in vitro* metrics were associated with lesion outcomes within each baseline severity class. We therefore analyzed severe and less severe lesions separately. Because *in vitro* combination metrics predict *in vivo* outcomes in simpler pathology models (Larkins-Ford et al., 2021, 2022), we expected equipotent assays in RMM-validated conditions to align more closely with less severe cellular/fibrotic lesions, whereas LIDs and lesion-informed dosing would align more closely with severe lesions.

To test these hypotheses, we stratified lesions by baseline severity and calculated Spearman rank correlations separately within each class, with each class’s correlations computed between *in vitro* metrics and the corresponding lesion outcomes (Data S1). We used the end-of-treatment measurements and the difference between the end-of-treatment and start-of-treatment measurements for both radiodensity and total lesion volume. We chose to examine end-of-treatment measurements to understand the final condition of marmosets within a single treatment arm, and we analyzed the difference between the end of treatment and the start to assess the degree of change. Given the limited number of combinations in our NHP dataset, we focused on trends and magnitudes of correlates to avoid overinterpretation. In the less severe lesions, we found very few significant correlations between *in vitro* metrics and lesion outcomes (Figure S2); moreover, the observed correlations were weak in magnitude.

To identify *in vitro* features associated with treatment response in clinically relevant disease, we focused subsequent analysis on severe lesions, as less severe lesions exhibited few and weak correlations with *in vitro* metrics (Figure S2). In severe lesions, we observed multiple significant associations between lesion-level PET/CT outcomes and *in vitro* drug-response metrics (Figure 3A).

**Figure 3.**
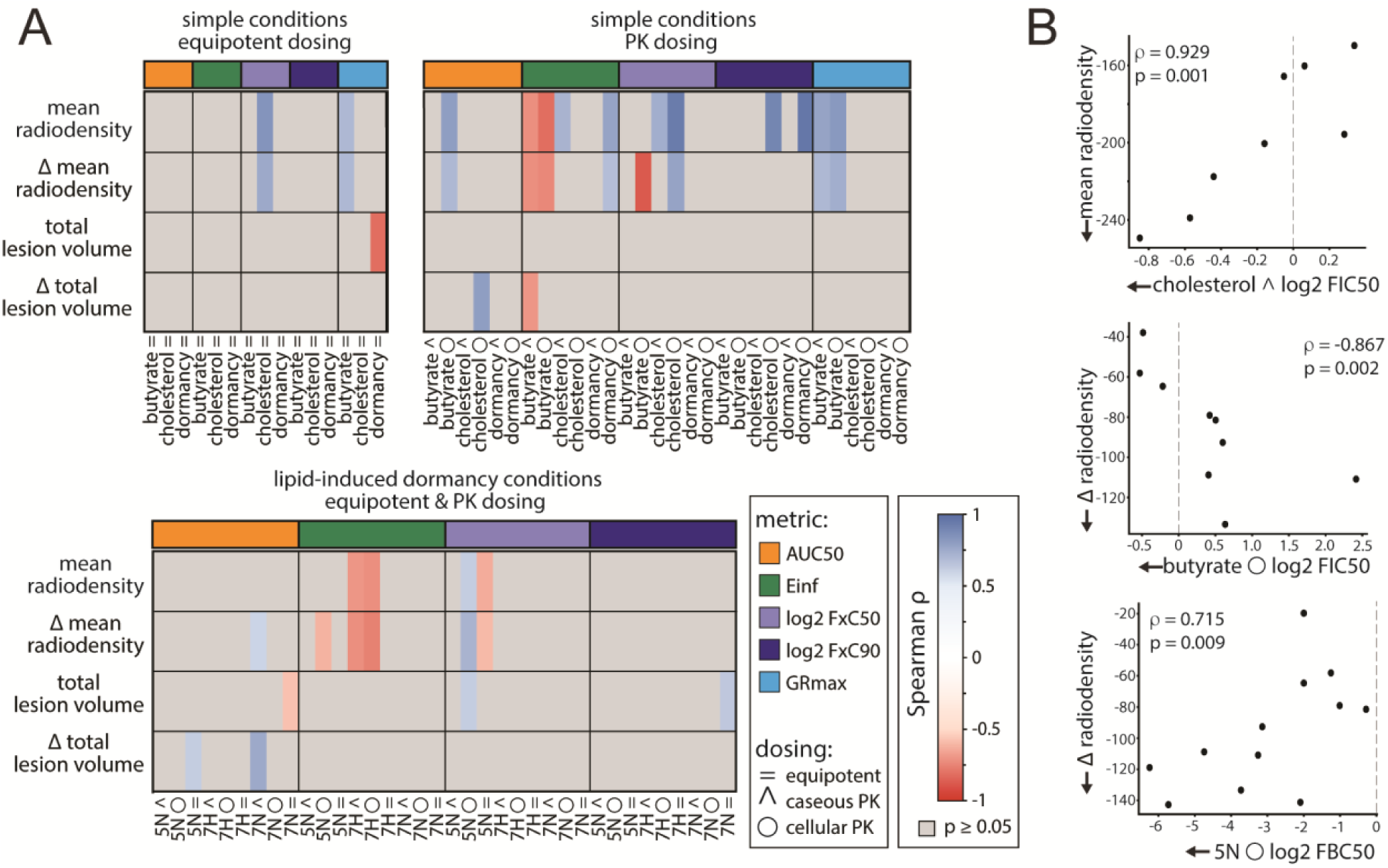
Correlate analysis for severe lesions with *in vitro* metrics. (A) Heatmap illustrating correlations between severe lesions and *in vitro* metrics. Equipotent dosing strategies in simple conditions have the lowest correlation frequency. Conditions designed to mimic the severe lesion environment (LIDs) and those with pharmacological dosing ratios show stronger correlations. No correlations with a p-value of greater than 0.05 are shown. (B) Scatter plots of select positive and negative correlations. The top panel illustrates an example of a correlation between cholesterol cellular PK log2 FIC50 of cholesterol and mean radiodensity. The middle plot shows the correlation between butyrate caseous PK log2 FIC50 and the delta in radiodensity. The bottom panel shows the correlation between acidic normoxic log2 FBC50.

In severe lesions, significant associations were concentrated in CT-derived radiodensity endpoints, whereas PET activity and volumetric measures showed few consistent associations. Across *in vitro* conditions, LIDs assays and PK-dosing (cellular and caseous) accounted for the most significant correlations, while simple equipotent assays contributed relatively few. This pattern indicates that lesion-mimicking growth conditions and lesion-relevant dosing ratios capture information that is more predictive of treatment response in severe pathology.

Some *in vitro* features were unavailable under dormancy conditions, specifically growth-rate-dependent metrics (e.g., GRmax), due to limited bacterial replication. Additionally, several high-kill metrics (FBC90) were undefined in many LIDs conditions because the bactericidal effect plateaued below 90% kill. Correlation patterns were similar for end-of-treatment radiodensity and its change from baseline, reflecting their inherent dependence. Lesions that exhibited larger reductions in radiodensity necessarily reached lower final radiodensity values. The same relationship was observed for lesion volume.

Representative examples of *in vitro* metric and pathology metric correlations are shown in Figure 3B. In cholesterol medium with cellular PK dosing, the interaction metric (log2 FIC50) was strongly negatively correlated with mean radiodensity at the end of treatment (ρ = −0.929, p = 0.001). Similarly, interaction metrics in butyrate under cellular PK dosing at FxC50 were strongly correlated with the magnitude of radiodensity decline over the course of treatment (ρ = −0.867, p = 0.002).

Together, these single-metric correlations indicate that lesion-relevant *in vitro* measurements contain signal for severe lesion treatment response, but no individual metric fully explains lesion trajectories. We therefore asked whether integrating these signals with longitudinal PET/CT measurements could improve the prediction of lesion progression.

### Prediction of marmoset lesion progression

To test whether these imaging and *in vitro* signals could be integrated to predict severe lesion trajectories, we developed computational models using random forest regression algorithms because of their ability to model multivariate, non-linear feature interactions in an interpretable manner. These models were tasked with predicting radiodensity at various time points. Models to predict PET activity were also tested; however, our models were unable to predict PET activity accurately and consistently (Figure S3). We anticipated this outcome because of the few correlates. We therefore attempted to predict radiodensity alone in severe lesions.

We developed three distinct computational models to investigate the hypothesis that lesion-level outcomes of severe lesions at the start of treatment can be predicted using data from marmoset PET/CT pathologies, with and without accompanying *in vitro* data. In the modeling analyses, we use the term *feature* to mean any model input variable, including PET/CT measurements, prior radiodensity values, treatment labels, and *in vitro* drug response metrics. The first model, referred to as the <u>naïve model</u>, used a naïve approach, benchmarking the information carried by the baseline lesion state alone. As an expected baseline comparator, we first asked how much of the end-of-treatment radiodensity could be explained by baseline PET/CT measurements alone. This naïve model captured only coarse persistence of lesion state, with an R² of 0.37, indicating that baseline pathology alone is insufficient to forecast treatment-modified lesion trajectories, highlighting the need for more informative predictive models.

In the second model type, referred to as <u>marmoset-only</u>, we adopted a sequential modeling approach: models were trained to predict each two-week radiodensity from prior observations. Predictions were then passed forward as inputs to the next time step. The input to the initial model was the same as the naïve model and included the radiodensity at the start of treatment for each lesion. The target output was changed to the next time point, as we anticipated that predicting a change in treatment for 2 weeks would be easier for the model than predicting 8 weeks. The model ensemble was sequential so that the output of the first model (which predicted week 2 radiodensity) was added to the input of the second model (which aimed to predict week 4 radiodensity). This sequentially chained strategy captured temporal lesion dynamics more accurately than the naïve model, yielding R² values ranging from 0.44 to 0.76 (Figure 4A, blue). Examples of lesion-level progression predicted by this model are shown at the top of Figure 4b for both MRZE and BPaL. Prediction performance tended to improve over time, with the model’s lowest error occurring in week 6. All time points had lower error than the initial prediction, suggesting that as the models see more time points, their prediction accuracy improves, ultimately yielding a close representation of the lesion-level trajectory.

**Figure 4.**
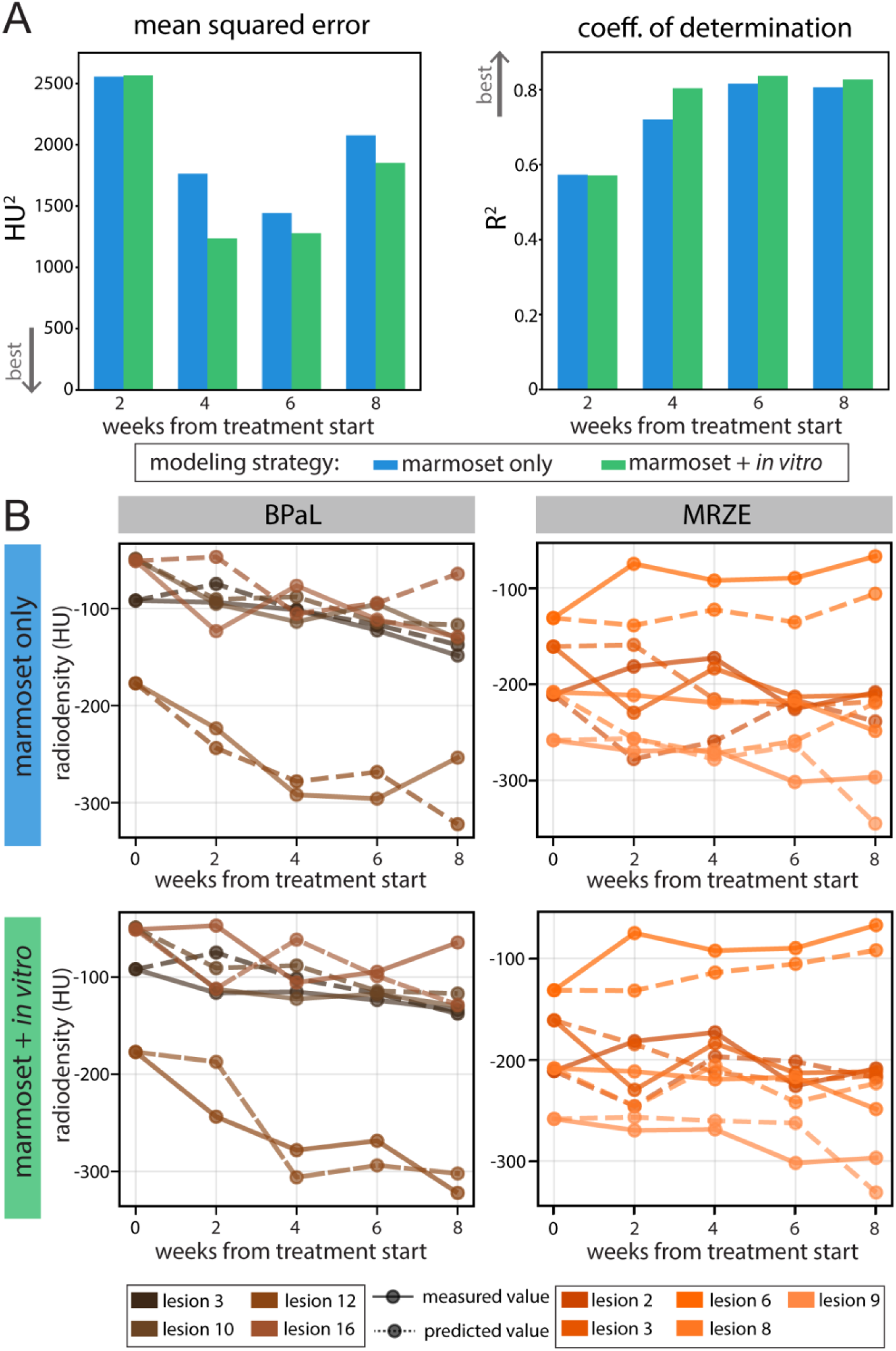
Predictive model performance and examples with sequential modeling. (A) Model performance metrics at the timepoints with available PET/CT data. Lower mean squared error (MSE) is considered better, and a higher coefficient of determination (R^2^) is considered better. The model trained on *in vitro* data performs better across nearly all time points and both metrics. (B) Lesion-level progression comparing actual lesion progression with model predictions.

In the third model type, referred to as <u>marmoset + *in vitro*</u>, we hypothesized that including corresponding *in vitro* treatment data would improve our ability to predict severe lesion progression over time. Using the same sequential framework, we augmented the model inputs with the treatment regimen administered to each lesion in the marmoset study. This addition moderately improved model performance on the held-out lesions (described in the Methods), increasing the range of R² from 0.58 to 0.85 (Figure 4A, green). In addition to improving R², this model also reduced the mean squared error (MSE) across all time points. Example predictions of lesion progression for BPaL and MRZE are shown at the bottom of Figure 4B, demonstrating the model’s ability to predict the impact of treatment on individual lesions. The improvement from adding *in vitro* data is slight; however, the improvement becomes more noticeable when comparing the distances between each week’s predictions for BPaL and MRZE. A clear example is the fine-tuning of predictions in the top lesion (6) of MRZE for both modeling types (Figure 4B), where the distance between the actual and predicted values is reduced for all *in vitro*-assisted predictions compared with marmoset-only predictions. The bulk of predictive power stems from sequential knowledge of the lesion over time, but fine-tuning is achieved by incorporating *in vitro* metrics, which bring the predictions closer to true values and ultimately improve performance across nearly every radiodensity output at each time point (Table S6).

### Relationship between in vitro drug response and predicted lesion outcome

We analyzed the radiodensity sequential model to understand how *in vitro* metrics predict the outcomes of marmoset lesions and to identify trends in high-performing combinations. SHapley Additive exPlanations (SHAP) analyses of the sequential radiodensity model revealed that the radiodensity at the previous time point was the dominant predictor across all forecasted time points. Among *in vitro* predictors, PK-informed dosing metrics (cellular/caseous PK) and lesion-mimicking LIDs features contributed most consistently, while equipotent RMM-validated condition metrics showed smaller, infrequent effects. A selected SHAP plot of the end-of-treatment radiodensity predictions is shown in Figure 5A, where the highest-impact features are the prior timepoint radiodensity measured, followed by *in vitro* features, predominantly those capturing PK-informed dosing and LIDs conditions. The spread of the data points in the SHAP plot corresponds to the magnitude of their impact on the model output, and the color of the data points is chosen so that blue indicates a lower feature value and red a higher one. Across time points, prior radiodensity is the most important predictor. LIDs-derived and PK-informed features follow in importance, indicating that baseline lesion state dominates, with lesion-relevant assay measurements providing additional signal.

**Figure 5.**
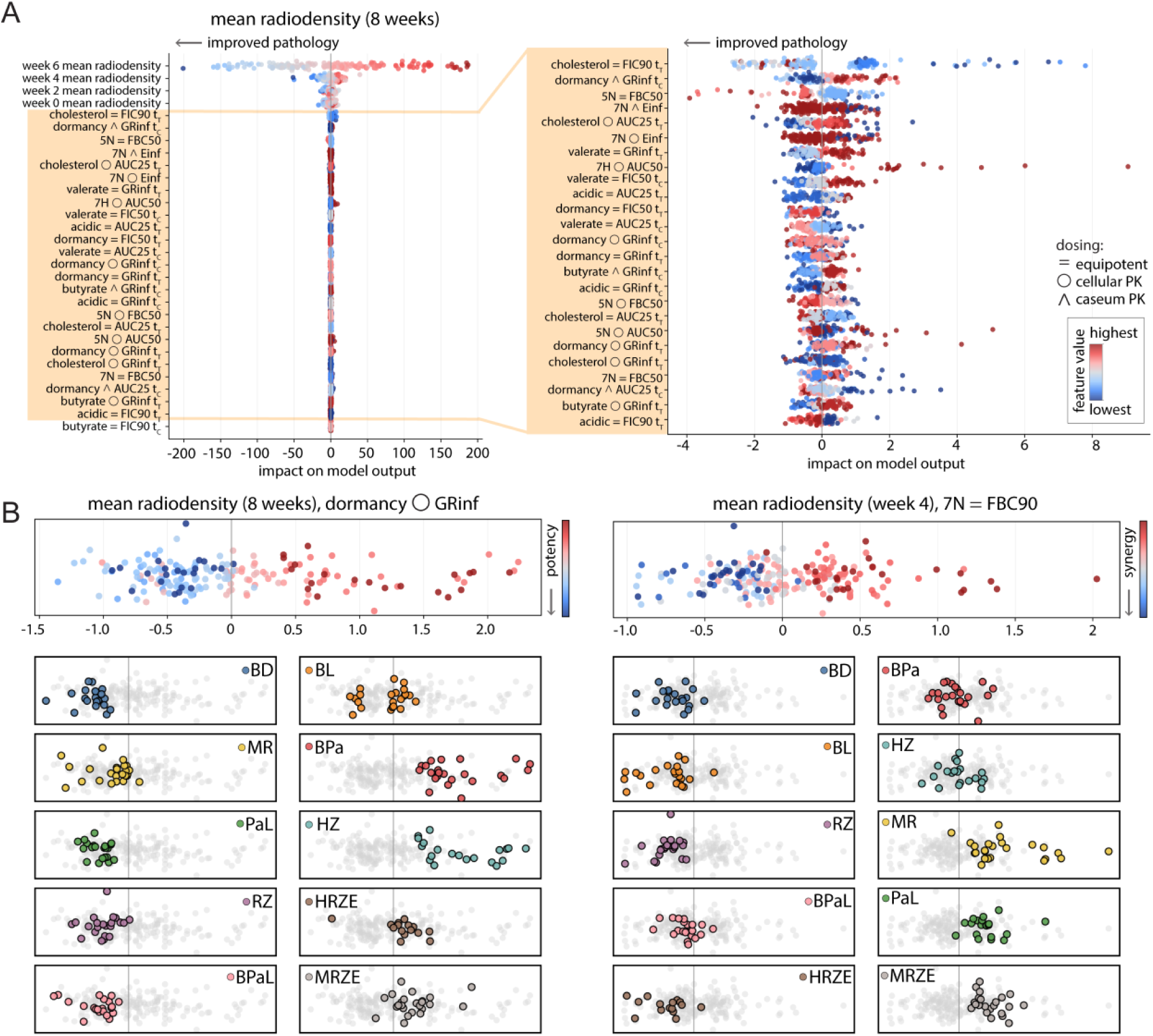
Relationship between *in vitro* metrics and sequential predictions of lesion radiodensity. (A) Global feature importance by SHAP for end-of-treatment radiodensity. Bee swarm plots of SHAP values from the sequential random forest model predicting week 8 mean radiodensity. Each point is one lesion. The X-axis is the SHAP impact on model output, where negative values mean a lower predicted radiodensity and thus improved pathology. The point color encodes the feature’s value, where blue represents a lower value. Prior radiodensity features, predicting weeks two to six, dominate the model’s influence, followed by *in vitro* metrics, most prominently those measured under PK-informed dosing and in LID conditions. (B) A zoomed-in view of the *in vitro* features. The same SHAP bee swarm is shown, but restricted to *in vitro* predictors, to visualize lower-magnitude effects. PK-informed metrics and LIDs features contribute more consistently than equipotent, non-LIDs metrics. (C) Interpretable examples with regimen stratification using individual SHAP features. Two features showing a clear value-to-effect bifurcation are presented, with the left panel illustrating contributions from cellular PK dosed GRinf in nitrate-induced dormancy at the end of treatment, and the right panel showing contributions to week four predictions from equipotent dosed neutral normoxic FBC90. Treatment regimens are colored individually in these feature-SHAP plots, showing groups in which there is a consensus of improvement predicted by the feature (indicated by being left of the dashed line, or contributing a negative value to the prediction).

To identify interpretable features, we sought features with a clear bifurcation between feature value and magnitude, such that improvement (to the left of the Y-axis in the SHAP plots) was associated with primarily red feature points and decline with blue. A magnified illustration of the spread of *in vitro* features is shown in Figure 5B, where the pathological features are cropped to display the spread of lower-magnitude metrics. We highlight two features that met this criterion in the top portion of Figure 5C, where the only SHAP rows shown are GRinf with caseous PK dosing in simple dormancy for week 8 radiodensity, and FBC90 with cellular PK dosing in neutral normoxic conditions for week 4 radiodensity. Lower feature values for these metrics are interpreted as more potent and more synergistic, respectively. In both cases, the lower feature values (in blue) are primarily on the left-hand side of the dashed line representing Y=0, indicating improvement in pathology. Coloring by treatment regimen (rather than feature value) revealed grouping patterns, allowing us to see which drug combinations were associated with improved pathology across lesions (Figure 5C, bottom). In predictions for week 8, these treatments were expected to improve pathology, based on the model’s learning about GRinf in nitrate-induced dormancy, with cellular PK dosing ratios: BD, BL, BPaL, PaL, MR, and RZ (Figure 5C, left). According to the model’s predicted contribution of FBC90 in neutral normoxic conditions with equipotent dosing ratios, BD, BL, RZ, BPaL, HRZE, and occasionally BPa and HZ were expected to decrease pathology (Figure 5C, right) in week 4. The data from the earlier time point suggest that the HRZE drug combination is beneficial when introduced early in the treatment course, but its effectiveness appears to diminish as treatment progresses, consistent with previous findings (Greenstein, 2026). Together, these SHAP analyses show that while prior radiodensity carries most predictive weight, lesion-mimicking *in vitro* features provide interpretable, regimen-specific contributions that refine radiodensity forecasts.

## Discussion

Our study demonstrates that integrating lesion-level imaging with *in vitro* drug response metrics can predict tuberculosis lesion outcomes in a lesion-type-dependent manner. Potency and interaction metrics from simple *in vitro* models (validated with the RMM) align with outcomes in less severe, cellular/fibrotic lesions, consistent with prior mouse work (Larkins-Ford, 2021; Larkins-Ford, 2022). By contrast, predicting responses in advanced (necrotic, caseous, cavitary) lesions required lesion-mimetic LIDs conditions and PK-informed dosing. This divergence highlights that the microenvironmental context fundamentally reshapes regimen performance, mirroring clinical observations that necrotic and cavitary diseases respond less favorably to standard therapy (Imperial, 2018; Malherbe, 2026). The relative scarcity of correlates in less severe lesions likely demonstrates that most standard regimens achieve adequate effects in these easier-to-treat phenotypes, thereby reducing the discriminative power of *in vitro* features.

Severe lesions are often lipid-rich and can be hypoxic and acidic, conditions that slow Mtb growth and increase tolerance to multiple antibiotics (Via, 2008; Harper, 2012; Baker, 2014; Sarathy, 2018). The LIDs assays were designed to reproduce these stresses using a defined lipid matrix with controlled pH and oxygenation. Across drugs, LIDs measurements recapitulated key features of caseum-associated tolerance, including elevated bactericidal thresholds and partial agreement with the cross-drug susceptibility ordering observed in *ex vivo* caseum (Sarathy, 2018; Sarathy, 2023). We therefore view LIDs as a scalable complement to established preclinical systems (BALB/c relapse, C3HeB/FeJ, NHP imaging, *ex vivo* caseum, and *in vitro* caseum surrogates) for prioritizing regimens against drug-tolerant lesion states (Harper, 2012; Li et al, 2015; Xu et al., 2019; Tasneen et al., 2022; Sarathy, 2023; Greenstein, 2026).

These findings align with recent *in vitro-to-in vivo* translation work by Goh et al., who showed that assay signatures reflecting host- and lesion-relevant bacterial states, including ex vivo caseum in chronic infection, improved the prediction of preclinical efficacy and early bactericidal activity (Goh, 2025). Our study extends this concept to combination drug responses and lesion-resolved NHP imaging, showing that lesion-mimicking assays provide additional predictive signal specifically in severe lesions.

To interpret the factors driving the sequential model’s forecast, we examined feature combinations across time points. As expected, the strongest predictor of a lesion’s radiodensity at the next scan was its radiodensity at the prior scan, reflecting persistent lesion-specific trajectories over treatment. *In vitro* features contributed smaller, but still interpretable changes, and metrics measured in lesion-mimicking conditions and under PK-informed dosing ratios tended to shift predictions toward lower radiodensity. SHAP patterns highlighted specific potency and interaction features associated with predicted pathology improvement. Regimen-level grouping in these plots suggests that the model learned consistent treatment-associated contributions across lesions. Interestingly, features linked to bedaquiline-containing regimens carried greater influence at later time points, aligning with bedaquiline’s delayed bactericidal activity. Conversely, we emphasize that our design does not separate drug penetration effects from intrinsic tolerance, as PK-informed dosing is simply a proxy for lesion exposure rather than a direct measurement (Prideaux, 2015; Ordonez, 2020; Strydom, 2019).

Although our study was not designed to test resistance and evolution directly, the observation that some antagonistic *in vitro* interactions tracked with better lesion outcomes is consistent with primary studies showing that suppressive or antagonistic drug combinations can select against resistance in other bacterial systems (Chait, 2007; Michel 2008). Together with earlier work centered around the RMM, these results argue against a narrow focus on synergy alone (Larkins-Ford, 2021; Larkins-Ford, 2022). Moderate antagonism *in vitro* may be acceptable, or even advantageous in complex lesions, depending on the regimen and *in vitro* experimental context. An important consideration is that the interaction scores measured in our *in vitro* assays reflect *in vitro* profiles; antagonism, even in lesion-like experimental environments, does not necessarily indicate that drug combinations are truly antagonistic *in vivo* or in the lesion microenvironment.

Severe lesion phenotypes (extensive caseation and cavitation) are consistently difficult to sterilize (Sarathy, 2018; Cicchese, 2020; Lanoix, 2015; Irwin, 2014). Our framework offers a practical approach to pressure-testing regimens under lesion-mimicking conditions before committing to complex, expensive animal studies or trials. Because extensive caseous/cavitary disease sets the sterilization bar that ultimately dictates treatment length, regimen development in tuberculosis prioritizes optimizing regimens for these severe phenotypes (Imperial, 2018). The goal is equitable shortening: shorter courses that remain effective for patients with hard-to-treat lesions, not only those with milder disease.

Although no single *in vitro* feature cleanly separated improving from non-improving lesions on its own, model performance improved when several modest concordant signals were considered together. Because the feature space is high-dimensional and most individual effects are modest, we interpret SHAP values in aggregate rather than as a singleton. Practically, this means we looked for aligned signals across time and assay types (such as dormancy potency and PK-informed synergy) and weighed them alongside the current lesion state: a few weak but consistent *in vitro* cues, when matched to the lesion’s baseline and early trajectory, can shift predictions towards improvement. These compact, time-aware rules, given a severe baseline, provide actionable outputs for screening regimens and selecting dosing ratios.

<u>This</u> study has several limitations. First, we modeled radiodensity trajectories, a composite tissue endpoint reflecting changes in necrosis, fibrosis, and aeration, rather than direct measures of viable bacterial burden or relapse risk. Moreover, early kill surrogates, such as early bactericidal activity, are themselves imperfect predictors of longer-term outcomes and treatment-shortening potential. Second, PK-informed dosing ratios were derived from simulated compartment exposures and therefore serve as proxies for lesion penetration rather than direct measurements (Prideaux, 2015; Ordonez, 2020; Strydom, 2019). True lesion PK can vary substantially across hosts and lesions. Third, limited sample sizes per regimen and the number of severe lesions constrain power and increase uncertainty in interaction and potency metrics. Fourth, under highly tolerant dormancy conditions, some high-kill metrics (FBC90) were undefined because bactericidal effects plateaued below 90% kill. We view this as an informative signature of extreme tolerance and as motivation to use complementary ensemble metrics rather than treating it as a shortcoming. Finally, models were trained on a single NHP study with a specific strain and infection protocol, and generalizability across additional cohort strains and dosing schedules remains to be demonstrated.

Future work should validate these findings in additional NHP datasets, expand the regimen set, and integrate complementary readouts (e.g., histopathology, immune profiling, and, where feasible, microbiological endpoints) to strengthen the connection between imaging trajectories and treatment success outcomes. Future experimental work should focus expanding lesion-mimicking condition panels and coupling them to systematically curated *in vivo* datasets should further improve model performance and sharpen severity-aware regimen prioritization.

Our work demonstrates that combining lesion-mimicking *in vitro* assays with lesion-level imaging advances TB regimen design toward severity-aware, shorter, and more effective treatment strategies, particularly for lesion types that have historically been the hardest to cure.

## Materials and Methods

### Animals, ethics approval

All procedures with *C. jacchus* (common marmosets), including breeding in National Institutes of Health (NIH) facilities, were in accordance with the recommendations of the Guide for the Care and Use of Laboratory Animals of the NIH and approved by the National Institute of Allergy and Infectious Diseases Animal Care and Use Committee in protocol LCIM-9 (permit issued to NIH as A-4149-01). Both male and female marmosets, ages 2 to 6 years, were included in the study, but as group numbers were small, no sex-specific analyses were conducted. Efforts were made to minimize suffering and provide intellectual and physical enrichment as approved in LCIM-9. Further details on the experimental design of the marmoset studies can be found in previous work (Greenstein, 2026).

### Marmoset PET/CT imaging, lesion tracking, and feature extraction

Treatment began 6 to 7 weeks after infection. PET/CT scans were performed before treatment, at treatment start, and after 2, 4, 6, and 8 weeks of treatment. At week 8, animals were sacrificed, and PET/CT image-guided necropsy was performed for lesion collection and CFU enumeration. To track individual lesions longitudinally, PET/CT scans were co-registered across time points. Regions of interest (ROIs) were demarcated from CT images based on radiodensity measured in Hounsfield units (HU). Structures with radiodensities greater than −500 HU were classified as lesions (Coleman, 2014; Chen, 2014). Voxels with radiodensities between −500 and −100 HU were designated as soft tissue, and voxels between −100 and 200 HU were designated as hard tissue. Voxel counts were converted to volumes within each ROI to quantify soft tissue volume, hard tissue volume, and total lesion volume. PET activity was quantified within each CT-defined ROI as FDG standardized uptake value (SUV). These extracted features generated longitudinal lesion-level measurements of PET activity, radiodensity, and lesion volume for subsequent severity classification, correlation analysis, and predictive modeling. Additional details are described in Greenstein (2026).

### Bacterial culture and media

All Mtb experiments were performed with Mtb Erdman transformed with pMV306hsp+LuxG13 to generate an autoluminescent strain (Addgene plasmid #26161, http://n2t.net/addgene:26161, RRID: Addgene_26161) as previously described (Larkins-Ford et al., 2021). Mtb was cultured in 7H9 Middlebrook medium supplemented with 0.2% glycerol, 10% ADC (50g/L albumin, 20g/L dextrose, and 0.04g/L catalase), and 0.05% Tween-80 with 25 μg/mL kanamycin (standard media) at 37 °C with aeration and shaking unless noted. Mtb stocks were frozen in media with 25% glycerol. Cells were passaged at mid-log phase (between OD600 = 0.5 – 0.8)

### RMM-validated *in vitro* conditions

Equipotent measurements in RMM-validated *in vitro* conditions were performed as described previously (Larkins-Ford et al., 2021; Larkins-Ford et al., 2022). For PK-informed measurements in the same media conditions, the culture conditions and assay endpoints followed these prior protocols, but drug concentrations and combination ratios were re-centered using the cellular or caseous lesion C_max_ values described below.

### LIDs media preparation

Base media (both acidic and neutral) was composed of 4.7g/L 7H9 powder, 0.5g/L fatty acid-free BSA, 100 mM NaCl, and 0.05% tyloxapol in deionized water. Neutral base media was buffered to pH 7.0 with 100 mM 3-(N-morpholino)propanesulfonic acid (MOPS). Acidic base media was buffered to pH 5.5 with 100 mM 2-(N-morpholino)ethanesulfonic acid (MES). Lipids were dissolved in a 1:1 (v/v) mixture of ethanol and tyloxapol that was pre-warmed to 80 °C for 30 minutes and added to pre-warmed (37 °C) base media at the following final concentrations: 150 μM cholesterol, 50 μM palmitic acid, 45 μM stearic acid, 4 μM oleic acid, 10 μM linoleic acid, and 3 μM arachidonic acid. The neutral-hypoxic (7H) media were also supplemented with 5 mM sodium nitrate. The pH was checked and adjusted with sodium hydroxide as needed after the lipids were added and all media were filter-sterilized using a 0.2 μm filter.

### LIDs acclimation and assay preparation

Mtb was thawed in standard media and sub-cultured at mid-log phase twice prior to use in LIDs assays. LIDs media was made fresh for each assay 1 day prior to use and maintained at 37 °C to prevent lipid precipitation. For acclimation, Mtb was grown to mid-log phase and resuspended in LIDs media at a concentration of OD600 = 0.5. The neutral-normoxic and acidic-normoxic conditions were incubated with aeration and shaking for 4 weeks at 37 °C. The neutral-hypoxic condition was incubated without aeration or shaking for 4 weeks at 37 °C. After 4 weeks of acclimation, Mtb cultures were syringe-passed through a 25-gauge needle twice to achieve a homogenous (non-clumpy) suspension for plating in 384-well plates.

### Charcoal agar plates

7H10 Middlebrook agar plates were supplemented with 0.4% activated charcoal, 0.5% glycerol, 10% ADC, 0.05% Tween-80, and 25 μg/mL kanamycin. These plates must be poured in a biosafety cabinet under sterile conditions to prevent contamination. 200 μL of 7H10 media was dispensed per well in 96-well plates using a multichannel pipette. Tips must be changed frequently to prevent clogging. Media was kept warm (low heat setting so as not to burn) on a heated plate with continuous stirring to prevent precipitation of charcoal throughout dispensing. Plates were poured within 7 days of use and stored at 4 °C until the day before use. They were then stored at room temperature for up to 24 hours under sterile conditions prior to use.

### Recovery and plate measurements

Dormant Mtb are characterized by low metabolic activity and therefore express low luminescence levels after 4 weeks of acclimation to dormancy (LIDs) media (Figure S1). To measure drug response in dormant bacilli, we measured regrowth upon recovery after treatment. For recovery of dormant cells, a solution of sterile PBS supplemented with 0.2% tyloxapol (PBS-Tx) was prepared. 40 μL of PBS-Tx was plated in all wells. The cells were mixed by pipetting, and 30 μL of cells/well were transferred to charcoal agar in 96-well plates. Plates were left to dry in the hood for several hours (no longer than 4 to prevent overdrying) and then incubated at 37 °C. The neutral-normoxic condition was incubated for 4 days, acidic-normoxic for 6 days, and the neutral-hypoxic for 8 days. Luminescence measurements were made using a Synergy Neo2 Hybrid Multi-Mode reader. Times were chosen based on detectable recovery of luminescence. Failure to regrow is indicative of cell death due to drug treatment, and therefore, the functional outcome of the dose-response curves was percent kill.

### Drug treatment

The drugs used in this study are listed in Table 1. All drugs were reconstituted in DMSO. Drugs were dispensed randomly using an HP D300e digital dispenser so that the final DMSO concentration per well did not exceed 1%. Drugs were dispensed at a 10-dose resolution with 2-fold spacing. For each LIDs condition, 40 μL of cells were plated per well in 384-well plates with freshly dispensed drugs, and plates were incubated at 37 °C for 7 days. The neutral-hypoxic condition plates were sealed with PCR seals to reduce oxygen exposure. Each plate contained multiple untreated and positive control wells for quality control.

### Dose selection for equipotent dosing and lesion-pharmacokinetic dosing

For the equipotent experiments, dose-centering experiments were performed for each LIDs condition as previously described (Larkins-Ford et al, 2021). In brief, the concentration to achieve 90% kill (BC90, determined as failure to regrow to 10% of untreated levels) for each single drug was tested. The BC90 was then used as the 7th dose out of 10 in combination measurements, where doses were 2-fold spaced to capture the treatment’s full range of response. If BC90 was not reached, the dose was centered around the C_max_ and/or DMSO solubility limits. For lesion-centric pharmacokinetic dosing, C_maxes_ in caseum and cellular lesions were determined for bedaquiline, linezolid, pretomanid, isoniazid, rifampicin, ethambutol, pyrazinamide, and moxifloxacin using GranSim (Methods). C_maxes_ for the remaining drugs in caseum and cellular lesions were determined or approximated from communications with VAD or published literature, where available.

### C_max_ determination for simulation of lesion pharmacokinetics

GranSim, a multi-scale, agent-based computational model of Mtb host-pathogen interactions, was used to determine the maximal achievable concentration (C_max_) of specified antibiotics in caseous and cellular lesions. GranSim simulates agents (e.g., macrophages, T cells, Mtb, chemokines, cytokines) in a 6mm x 6mm area of lung tissue represented by a grid structure. The cellular/molecular processes and interactions between the agents are defined by parameterized mathematical equations based on the immunological rules from nonhuman primate (NHP) data (Cilfone et al., 2015; Hult, Mattila, Gideon, Linderman, & Kirschner, 2021; Marino et al., 2015; Ray, Flynn, & Kirschner, 2009; Sud, Bigbee, Flynn, & Kirschner, 2006; Warsinske, DiFazio, Linderman, Flynn, & Kirschner, 2017). Granulomas are grouped based on colony-forming unit (CFU) trends into two categories: high and low CFU granulomas (Budak et al., 2023). High CFU granulomas are caseous granulomas with uncontrolled bacterial growth, whereas low CFU granulomas have less caseum and are composed mostly of immune cells with stable bacterial CFU levels (Budak et al., 2023). A simulated granuloma library of 200 granulomas (100 high CFU and 100 low CFU) was created for C_max_ calculations.

The concentration of antibiotics within simulated granulomas is modeled based on the pharmacokinetic (PK) processes occurring within granulomas. Antibiotics permeate into the lungs via vascular sources, which is simulated in the GranSim environment (Cicchese, Dartois, Kirschner, & Linderman, 2020; Cicchese, Sambarey, Kirschner, Linderman, & Chandrasekaran, 2021; Pienaar, Cilfone, et al., 2015; Pienaar, Dartois, Linderman, & Kirschner, 2015; Pienaar et al., 2017). Once on the grid, antibiotics diffuse, bind to macromolecules, and are taken up by immune cells (Cicchese et al., 2020; Cicchese et al., 2021; Pienaar, Cilfone, et al., 2015; Pienaar, Dartois, et al., 2015; Pienaar et al., 2017). Each of these processes is defined by mathematical equations whose parameters are calibrated to in vivo studies from human or animal models (Budak et al., 2023). GranSim simulates the PK of eight different antibiotics: isoniazid, rifampicin, pyrazinamide, ethambutol, moxifloxacin, bedaquiline, linezolid, and pretomanid.

To determine the average C_max_ of each of the 8 antibiotics in caseous and cellular lesions, treatment of the generated granuloma library is simulated using human or human-equivalent dosing for 24 hours (Budak et al., 2023): 5 mg/kg of INH, 10 mg/kg of RIF, 7mg/kg of MOX, 20 mg/kg of EMB, 25 mg/kg of PZA, 20mg/kg of BDQ, 20mg/kg of PRE and 90 mg/kg of LIN. Antibiotic concentrations in caseum of high-CFU granulomas at each time step (C_cas_(t)) were tracked to calculate the average C_max_ in caseum. Antibiotic concentrations in low-CFU granulomas at each time step (C_cell_ (t)) were tracked to calculate the average C_max_ in cellular lesions. The average C_max_ in cellular and caseous lesions was calculated using the following equations:

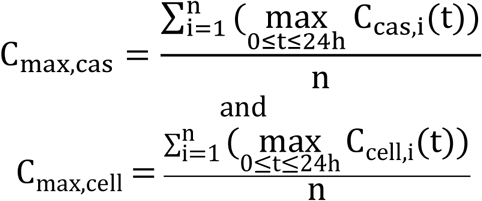

where *n* is the number of granulomas, *t* is the simulation time, *C_cas_* and *C_cell_* are average antibiotic concentrations of granuloma *i* from low-CFU granuloma set and the caseum of high-CFU granuloma set, respectively.

### Hill fitting and quality control

After normalizing drug response data to untreated and inverting (subtracting from 1), values greater than 1 (=100% kill) were set to 1, under the rationale that there is no biological interpretation to >100% kill. Values less than 0 were set to 0 by the rationale that there is no biological interpretation to <0% kill. These data exhibited greater variability than simpler in vitro assays, and therefore, most fitted curves were visually inspected to assess the trustworthiness of the fit. The visual inspections were blinded and performed twice (by two different individuals) to prevent bias.

If the percent kill attributed to the highest dose of a drug or combination was <=0, the treatment was flagged “bad” and not fitted to a Hill curve with very few exceptions (e.g., strong reason to suspect drug precipitation at the highest dose). Dose-response curves that did not exceed 50% kill with monotonic increase were flagged “bad” and not fitted to a Hill curve. Dose-response curve data in which only the last 1-3 points achieved greater than 50% kill (i.e., only the last 1-3 points were deemed trustworthy) were fitted using 0s for the first 7-9 points and the last data point(s). Treatments flagged “bad” were assigned Hill slopes of 0. If the normalized percent kill for the highest dose of a “bad” treatment was <0, the assigned Einf was 0; if it was between 0 and 50, the assigned Einf was 0.25 (25%).

If the percent kill attributed to the highest dose dipped more than 10% lower than the previous dose, we assumed precipitation at the highest dose, discarded the last point, and flagged the fit for visual inspection to check if the assessment of precipitation appeared reasonable. Data points preceding dips in percent kill that exceeded 40% were considered too variable to trust and discarded. Fits with data points preceding dips in percent kill that exceeded 20% were flagged for visual inspection; if the dip was determined to affect the fit, the data preceding the dip were discarded. As appropriate, individual data points were deleted if they were visually determined to be outliers (e.g., a substantial dip in kill after multiple points > 80% kill and multiple points > 80% kill following the dip) or otherwise evaluated to be untrustworthy and compromising the fit of the Hill curve. Fits that resulted in inappropriately low upper asymptotes (Einf) where the treatment clearly achieved a higher percent kill were refit with the last two data points weighted 10 times more than the rest to force a more accurate Einf.

### *In vitro* data processing

*In vitro* data processing and dose-response metric calculation were performed using custom MATLAB scripts. The median luminescence of empty (background) charcoal wells was subtracted from the raw data, and drug-treated wells were normalized to the 25th percentile of untreated wells within each plate. Dose-response data were fit to a three-parameter Hill function using the “NonlinearLeastSquares” method, and bactericidal concentrations were calculated from the Hill curve as previously described (insert cell systems ref). Dose-response curves that did not exceed 50% kill with monotonic increase were flagged “bad” and not fit to a Hill curve. Specific details of curve-fitting and quality control are expanded in Supplementary Methods. The area under the curve at 50% kill (AUC50 was calculated using the integral of the fit curves from 0 to the 50% kill concentration (BC50) and normalized to the BC50. The Einf (upper asymptote of the Hill curve) describes the maximum achievable effect by a given drug treatment. Drug interaction scores were calculated using the principles of Bliss Independence (details in supplementary materials).

### Clustering analysis of marmoset lesions

Details of the marmoset study design are provided in the Supplementary Materials. Two animals and one isolated lesion from a third animal developed repeated phenotypic resistance; these animals/lesions were removed from subsequent analysis (Table S7). Lesions without valid colony-forming unit (CFU) counts were removed. Data from the scan at the start of treatment was normalized and scaled using the standard scaler from Python sklearn. Leiden community detection was performed using the Python package scanpy. In brief, data was transformed using PCA, and all principal components were used to build a nearest neighbors graph (using cosine distance metric). Leiden community detection (LCD) (resolution = 1.0) was used to detect dense communities in the neighbor graphs. The Python package dashbio performed hierarchical clustering with complete linkage and cosine distance. “High severity” and “low severity” lesion clusters were determined based on the branching of the clustergram.

### Correlation analysis

Analyses were implemented in Python (primarily using pandas, scipy, and matplotlib) using two sources: (1) in vitro combination metrics and (2) PET/CT lesion-level data. Lesions were stratified by baseline severity before calculating the correlations. From the PET/CT dataset, we used the end-of-treatment radiodensity (i.e., radiodensity at timepoint 6, 8 weeks after treatment), the end-of-treatment total lesion volume (calculated as the sum of soft and hard volumes), and deltas, which were computed as the end-of-treatment value minus the start-of-treatment value. We considered metrics across all our assays and dosing schemes, including FxC50, FxC90, AUC50, Einf, and GRmax. We chose three non-LIDs conditions: butyrate, cholesterol, and nitrate-induced dormancy (referred to as “dormancy” in the figures). The three LIDs conditions designed were also included in the correlations. Dosing for each condition was performed at equipotent, cell PK, and caseous PK dosing. For each in vitro feature pathology pair within a severity class, we computed Spearman’s rank correlation (ρ) and two-sided p using the scipy package (scipy.stats.spearmanr). Given multiple lesions per drug combination, pathology values were aggregated to the drug-combination level by mean before merging with the corresponding in vitro metric. For the pathological features analyzed, a lower value is considered better. One *in* vitro feature, Einf, behaves in the opposite manner, such that a higher value in either of these two metrics corresponds to a more potent combination. Thus, to determine whether ideal combinations are high potency, we must look for negative correlations. The remaining features shown in Figure 3 exhibit the same directionality as the pathological data: lower values correspond to more synergistic or potent conditions. We report the direction and magnitude of ρ and display only significant pairs at α = 0.05. Heatmaps were rendered with a diverging colormap centered at zero, and non-significant pairs are plotted as light grey squares. Axes labels are grouped by their metric family (colored bar), and the condition in which they were measured in. They are additionally stratified by dosing strategy and LIDs assays. For the three example scatter plots in Figure 3B, we subset the severe cohort, aggregated pathology to the drug-combination level (mean), and plotted the chosen in vitro metric (X-axis) versus the corresponding pathological feature (Y-axis). The Spearman ρ and p are annotated in the panels.

### Modeling data preparation

Input features came from two different sources: (1) baseline lesion characteristics measured at timepoint 2 (start of treatment), including mean Hounsfield units (mean HU), and (2) in vitro metrics for assorted combinations, tested in the discussed growth/dormancy conditions. Missing measurements are handled natively by scikit-learn’s RandomForestRegressor. Quabodepistat-containing compounds are excluded from modeling due to the large amount of missing data.

### Sequential model architecture

For each target time point (2 weeks of treatment, through 8 weeks of treatment), we trained a separate random forest regressor. The feature set expanded sequentially. Models predicting week 2 used only baseline features (start of treatment and +/- in vitro measurements matching the treatment the marmoset received, if the marmoset was part of the + in vitro model). In contrast, models for subsequent timepoints incorporated both baseline features and predicted radiodensity values from all preceding timepoints. For example, the model predicting week 8 includes the predictions for weeks 2, 4, and 6. Each random forest in the ensemble is set up using the default parameters of scikit-learn and was seeded for reproducibility.

### Train-test split strategy

To ensure rigorous evaluation and prevent data leakage, we implemented a compound-stratified train-test split. Rather than randomly splitting individual lesions, we split the data at the compound level, ensuring that all lesions from a given drug combination were assigned exclusively to either the training set (80%) or the testing set (20%). Within each compound group, lesions were randomly (seeded) allocated to maintain the 80-20 split. This strategy enables the model to learn how to better represent more drug responses, as there are only ten in the training set, and removing them (which is very possible via random splitting) would be overly punishing.

### Model evaluation

Model performance was assessed on the held-out test set using three metrics: mean squared error (MSE), mean absolute error (MAE), and coefficient of determination (R2). These metrics were calculated by comparing predicted mean radiodensities with ground-truth measurements at each time point. MAE values are not shown directly in the manuscript but are easily accessible via GitHub.

### Feature importance analysis

We employed three methods to assess feature importance: (1) random forest mean decreases in impurity (MDI), (2) permutation importance calculated on the test set, and (3) SHAP (SHapley Additive exPlanations) values. The general trends of each of these outputs were the same, so SHAP analysis was focused on due to more expansive tooling. SHAP analysis was performed on a random sample of up to 1,000 training observations per model using TreeExplainer, providing both global feature importance rankings and sample-specific feature contribution profiles.

### SHAP analysis

To quantify the contribution of individual features to model predictions, we employed SHAP analysis using the TreeExplainer algorithm, which is optimized for random forest models. SHAP values provide a unified measure of feature importance by computing each feature’s contribution to a prediction across a dataset. For each trained timepoint model, we calculated SHAP values on a random sample of up to 1,000 training instances. The TreeExplainer algorithm computes exact Shapley values by leveraging the tree structure of the random forest.

We then created SHAP summary plots to visualize both the magnitude and direction of feature effects across all samples. In these plots, each point represents a single lesion, with the X-axis indicating the SHAP value (the impact on predicted mean radiodensity for a timepoint), and the Y-axis showing features ranked by mean absolute SHAP value (essentially, rank ordering), with the color representing normalized feature values (low to high). This visualization enables the identification of features with consistent directional effects versus those with context-dependent impacts.

To facilitate interpretation, we generated two versions of the summary plots: (1) a comprehensive plot displaying the top 30 features by mean absolute SHAP values, and (2) a “zoomed” version of the SHAP plot, which excludes all prior timepoint radiodensity values, as they heavily widened the axes.

To assess whether in vitro features collectively predicted favorable or unfavorable treatment outcomes for each drug combination, we selected individual features and, for each measurement (lesion), colored them by combination. This allowed us to identify trends in contributing a negative amount to the prediction (improvement) or a positive value (worsening).

## Author Contributions

Conceptualization: JJW, TG, JS, LEV, CEB, VD, DK, BA

Data Curation: JJW, TG, MPM, MB, LEV

Formal Analysis: JJW, TG, MB

Funding Acquisition: BA, DK, VD, CEB

Investigation: TG, DMW, AA, JDF, FG, AW, MPM, MB

Methodology: all authors

Project Administration: BA, DK

Resources: MB, LEV, CEB, JS, DK, VD, BA

Software: JJW, MB

Supervision: BA, DK, CEB, LEV

Validation: JJW, TG, MPM

Visualization: JJW, TG, MPM

Writing – Original Draft: JJW, TG, MPM, BA

Writing – Review & Editing: all authors

## Acknowledgements

We thank Otsuka Pharmaceutical Co. for providing delamanid and quabodepistat. We thank members of the Aldridge lab for helpful discussions, including Hanna Clutterbuck-Cook. This work was supported in part by the Gates Foundation INV-027276 to BA, INV-012045 to CEB, and INV-040485 to CEB. The conclusions and opinions expressed in this work are those of the author(s) alone and shall not be attributed to the Foundation. Under the grant conditions of the Foundation, a Creative Commons Attribution 4.0 License has already been assigned to the Author Accepted Manuscript version that might arise from this submission. Please note that works submitted as a preprint have not undergone a peer review process.

Animal data reported in this publication were supported in part by the Division of Intramural Research (CEB and LEV). Research reported in this publication was supported by the Division of Intramural Research, National Institute of Allergy and Infectious Diseases (NIAID), National Institutes of Health (NIH) of the National Institutes of Health under award numbers R01AI50684 to BA, DK, and VD, UM1AI7179669 to BA, and the University of Michigan Center for Data-Driven Drug Development and Treatment Assessment (DK and MB). This research was supported in part by the Intramural Research Program of the National Institutes of Health (NIH). The contributions of the NIH author(s) are considered Works of the United States Government. The findings and conclusions presented in this paper are those of the author(s) and do not necessarily reflect the views of the NIH or the U.S. Department of Health and Human Services.

## Data and Materials Availability

### Materials Availability

N/A

### Data and Code Availability

Processed *in vitro* data and the scripts required to generate the figures have been deposited on Figshare, with DOI: 10.6084/m9.figshare.32257020.

All original code and figures have been deposited at https://github.com/joshwhiteley/marmoset-paper and are publicly available as of the date of publication.

