## Supplemental Material for "*In vitro* dormancy models improve ability to predict treatment response in severe marmoset tuberculosis lesions"

**Supplementary Materials**

Figures S1-S3

Tables S1-S6

Box S1

**Supplemental Figures**

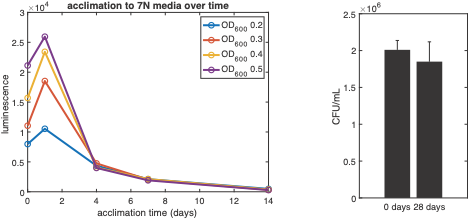

**Figure S1. Complex lipid media induce decreased metabolic activity indicative of non-replication.** Mtb were acclimated to the 7N condition for 28 days. Luminescence measurements were taken at days 0 (i.e., pre-acclimation), 1, 4, 7, and 14 of acclimation, and bacterial burden (colony-forming units, CFU) was enumerated at day 0. Luminescence readings were discontinued after 14 days because the luminescence decreased to the minimal unit of detection. Bacterial burden was enumerated again after 28 days of acclimation (full acclimation). No significant difference was determined between bacterial burden at days 0 and 28 (Student’s t-test, p > 0.05)

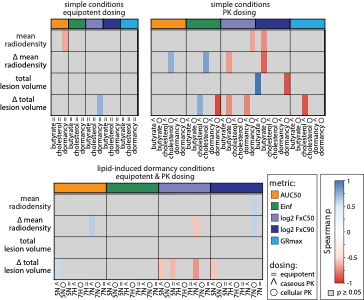

Figure S2. Correlate analysis for less severe lesions with *in vitro* metrics. An overall lower quantity of correlates are present in the less severe lesions than found in the severe lesions (Figure 3).

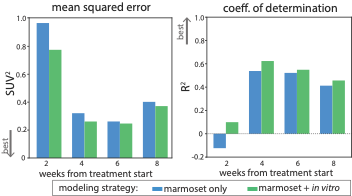

**Figure S3. Model performance metrics for a PET activity sequential model.** MSE and R2 for a sequential model with mean PET activity as the target for prediction. These models follow the same trends as the radiodensity only models (increase of performance with addition of in vitro data, and performance over time), however, are overall lower performing than the radiodensity model.

**Table S1. Median values and standard deviation for each feature within each cluster are described in Figure 2.**

|  | 0 | 1 | 2 | 3 | 4 | 5 | 6 | 7 | 8 | 9 | 10 | 11 | 12 | 13 | 14 | 15 |
| --- | --- | --- | --- | --- | --- | --- | --- | --- | --- | --- | --- | --- | --- | --- | --- | --- |
| PET activity (SUV) | 3.0 ± 0.9 | 1.9 ± 0.5 | 3.3 ± 1.1 | 1.2 ± 0.4 | 2.1 ± 0.7 | 1.8 ± 0.6 | 1.5 ± 0.3 | 0.7 ± 0.4 | 0.8 ± 0.3 | 1.1 ± 0.3 | 2.3 ± 0.4 | 3.0 ± 1.2 | 3.4 ± 0.7 | 1.2 ± 0.5 | 1.7 ± 0.4 | 2.4 ± 0.4 |
| radiodensity (HU) | -84.3 ± 41.0 | -246.6 ± 108.0 | -109.2 ± 60.7 | -239.1 ± 41.5 | -88.8 ± 51.3 | -179.9 ± 36.3 | -152.0 ± 28.7 | -405.1 ± 90.8 | -298.6 ± 34.1 | -219.6 ± 22.6 | -202.7 ± 47.4 | -149.5 ± 47.7 | -217.2 ± 77.7 | -323.3 ± 65.3 | -69.0 ± 37.7 | -89.9 ± 32.0 |
| hard volume (mL) | 0.76 ± 0.58 | 0.03 ± 0.03 | 0.16 ± 0.12 | 0.03 ± 0.03 | 0.13 ± 0.08 | 0.29 ± 0.24 | 0.06 ± 0.05 | 0.00 ± 0.00 | 0.00 ± 0.01 | 0.02 ± 0.02 | 0.04 ± 0.02 | 0.33 ± 0.25 | 0.10 ± 0.05 | 0.01 ± 0.01 | 0.43 ± 0.17 | 0.14 ± 0.08 |
| soft volume (mL) | 0.18 ± 0.12 | 0.04 ± 0.03 | 0.06 ± 0.03 | 0.05 ± 0.04 | 0.04 ± 0.03 | 0.21 ± 0.13 | 0.04 ± 0.03 | 0.01 ± 0.01 | 0.01 ± 0.02 | 0.03 ± 0.02 | 0.04 ± 0.02 | 0.14 ± 0.06 | 0.07 ± 0.04 | 0.02 ± 0.02 | 0.10 ± 0.04 | 0.05 ± 0.03 |
| std. dev. radiodensity (HU) | 156.8 ± 23.4 | 235.4 ± 29.4 | 166.5 ± 36.9 | 148.8 ± 16.5 | 137.5 ± 31.9 | 168.9 ± 14.7 | 174.2 ± 13.2 | 118.7 ± 25.1 | 156.5 ± 12.8 | 170.3 ± 9.3 | 164.9 ± 14.9 | 226.8 ± 35.4 | 243.9 ± 38.7 | 205.2 ± 17.4 | 139.7 ± 22.0 | 210.3 ± 14.5 |

**Table S2. MBC90 and IC90 values for a suite of *in vitro* models.** Bolded text indicates the highest concentrations tested (due to solubility or based on C_max_).

| drug | inhibition assays IC90 (µM) | | | | bactericidal assays MBC90 (µM) | | | | |
| --- | --- | --- | --- | --- | --- | --- | --- | --- | --- |
|  | acidic | butyrate | cholesterol | dormancy | 5N equip. | 7N equip. | 7H equip. | caseum surrogate | *ex vivo* caseum |
| ethambutol | 229 | 41.1 | 25.9 | 135 | **489** | **489** | **489** | 90 | **512** |
| quabodepistat | 0.00206 | 0.00124 | 0.00169 | 0.0219 | 3.1 | 4.6 | 5.7 | ND | **512** |
| isoniazid | 0.292 | 0.146 | 0.219 | 5.32 | 160 | 46 | 49 | ND | **512** |
| pretomanid | 0.557 | 0.752 | 0.139 | 4.98 | 1.8 | 5.2 | 0.92 | 62.5 | 70 |
| delamanid | 0.262 | 0.131 | 0.0187 | 5.37 | 1.9 | 4.7 | 1.1 | ND | 512 |
| SQ109 | 6.5 | 2.51 | 1.45 | 45.4 | **182** | 11 | 6.8 | 200 | 260 |
| linezolid | 1.33 | 4.3 | 3.17 | 7.86 | 296 | 310 | 78 | 512 | **512** |
| sutezolid | 0.736 | 0.226 | 0.198 | 2.12 | **42** | 6.8 | 5.1 | 8.0 | 16.0 |
| bedaquiline | 0.432 | 0.054 | 0.09 | 1.51 | 1.4 | 3.8 | 6.3 | 5.4 | 4.7 |
| clofazimine | 2.62 | 3.91 | 2.22 | 11.8 | 26 | 6.8 | 28 | **128** | **128** |
| rifampicin | 0.0486 | 0.0729 | 0.0851 | 0.0486 | 0.11 | 2.5 | 1.3 | 2.2 | 10.0 |
| rifapentine | 0.00255 | 0.251 | 0.319 | 0.16 | 0.094 | 0.34 | 0.64 | 2.5 | 10.0 |
| moxifloxacin | 0.772 | 1.2 | 0.722 | 0.897 | 7.5 | 4.7 | 2.8 | 1.5 | 0.6 |
| pyrazinamide | 7420 | 839 | 240 | **11900** | **812** | **812** | **812** | **600** | **8196** |

**Table S3. Spearman correlations between LIDs conditions and *ex vivo* caseum.**

|  | caseum MBC90 | 5N MBC90 | 7N MBC90 | 7H MBC90 |
| --- | --- | --- | --- | --- |
| caseum MBC90 | 1 |  |  |  |
| 5N MBC90 | 0.65 | 1 |  |  |
| 7H MBC90 | 0.57 | 0.82 | 1 |  |
| 7N MBC90 | 0.75 | 0.90 | 0.85 | 1 |

**Table S4. Intra-condition FxC90 correlation values.** Comparative correlations inside of each feature, with Spearman rho and p values displayed. *n* combinations represents the amount of combinations with available FxC90 data for correlate calculation.

| ***in vitro* condition** | **dose ratio 1** | **dose ratio 2** | **ρ** | p | ***n* combinations** |
| --- | --- | --- | --- | --- | --- |
| 5N | equipotent | caseous | 0.38 | 0.19 | 13 |
|  | equipotent | cellular | -0.31 | 0.21 | 18 |
|  | caseous | cellular | -0.27 | 0.40 | 12 |
| 7H | equipotent | caseous | 0.17 | 0.58 | 13 |
|  | equipotent | cellular | 0.14 | 0.65 | 13 |
|  | caseous | cellular | 0.52 | 0.07 | 13 |
| 7N | equipotent | caseous | -0.25 | 0.40 | 13 |
|  | equipotent | cellular | 0.21 | 0.48 | 13 |
|  | caseous | cellular | 0.32 | 0.29 | 13 |
| butyrate t_C_ | equipotent | caseous | 0.73 | 0.02 | 9 |
|  | equipotent | cellular | 0.52 | 0.15 | 9 |
|  | caseous | cellular | 0.83 | 0.01 | 9 |
| butyrate t_T_ | equipotent | caseous | 0.67 | 0.07 | 8 |
|  | equipotent | cellular | 0.50 | 0.21 | 8 |
|  | caseous | cellular | 0.58 | 0.10 | 9 |
| cholesterol t_C_ | equipotent | caseous | 0.69 | 0.06 | 8 |
|  | equipotent | cellular | 0.67 | 0.07 | 8 |
|  | caseous | cellular | 0.50 | 0.21 | 8 |
| cholesterol t_T_ | equipotent | caseous | 0.70 | 0.04 | 9 |
|  | equipotent | cellular | 0.62 | 0.08 | 9 |
|  | caseous | cellular | 0.12 | 0.77 | 9 |
| nitrate dormancy t_C_ | equipotent | caseous | 0.09 | 0.87 | 6 |
|  | equipotent | cellular | 0.26 | 0.62 | 6 |
|  | caseous | cellular | 0.80 | 0.01 | 9 |
| nitrate dormancy t_T_ | equipotent | caseous | 0.60 | 0.21 | 6 |
|  | equipotent | cellular | 0.60 | 0.21 | 6 |
|  | caseous | cellular | 0.71 | 0.11 | 6 |

**Table S5. C_max_ values for drugs based on GranSim.** *The cellular C_max_ for bedaquiline was >100ug/mL, which was not soluble in DMSO and >50x MBC90, which was deemed impractical to measure.

| **drug** | **caseum Cmax (ug/mL)** | **cellular Cmax (ug/mL)** |
| --- | --- | --- |
| bedaquiline | 12 | 25* |
| ethambutol | 22.5 | 28 |
| isoniazid | 0.5 | 0.6 |
| linezolid | 23 | 42 |
| moxifloxacin | 56 | 59 |
| pretomanid | 27 | 10 |
| pyrazinamide | 83 | 82 |
| rifampicin | 3.6 | 3.3 |

**Table S6. Model error metrics for the three modeling strategies tested for mean radiodensity predictions.**

| **model** | **time point** | **MSE** | **R^2^** |
| --- | --- | --- | --- |
| naïve | week 8 | 5079.7 | 0.37 |
| marmoset | week 2 | 3152.7 | 0.44 |
|  | week 4 | 2378.5 | 0.60 |
|  | week 6 | 1874.9 | 0.73 |
|  | week 8 | 2354.5 | 0.76 |
| marmoset + *in vitro* | week 2 | 2363.6 | 0.58 |
|  | week 4 | 1394.2 | 0.77 |
|  | week 6 | 1039.5 | 0.85 |
|  | week 8 | 2094.3 | 0.79 |

**Table S7. Marmoset lesions excluded from the study**. Highlighted rows indicate a marmoset that was deemed to develop resistance, and thus, all lesions were discarded from analysis.

| **marmoset ID** | **lesions excluded** |
| --- | --- |
| 750NB | 6 |
| B011 | 11 |
| BI01 | 14 |
| BI07 | 4, 12 |
| BI60 | 1 |
| BI76 | 1, 11 |
| BJ16 | 13 |
| BJ27 | 5, 12, 13 |
| BJ28 | 6 |
| BJ45 | 10 |
| BK21 | all |
| BM01 | 3 |
| BM07 | 21, 22, 24 |
| BM16 | 16, 20 |
| BM19 | 4 |
| BM27 | 7, 13 |
| BM28 | 13, 14 |
| BM34 | 9, 20, 23 |
| BM40 | 9, 18, 21 |
| BN05 | 15 |
| BN13 | 7 |
| BN14 | 5, 9, 10 |
| BN18 | 7, 9 |
| BN20 | all |
| BN22 | 9 |
| BN27 | 3 |
| BN36 | 12 |
| BO09 | 10, 12 |
| BO11 | 23 |
| BO22 | 8, 9 |
| BO23 | 6 |
| BO36 | 21, 22 |
| BO37 | 11, 13 |
| BO42 | 5 |
| BP02 | 21 |
| BP09 | 13 |
| BP14 | 11, 14, 27, 28, 29, 30, 44, 47, 50, 51 |
| BP17 | 4, 8, 12, 18, 26, 29 |
| BP20 | 7, 8, 12 |
| BP29 | 10, 15 |
| BP35 | 4, 15, 16 |
| BQ02 | 23 |
| BQ42 | 9, 13, 20, 22, 23 |
| BQ47 | 4, 10 |
| BR01 | 5 |
| BR07 | 12, 18 |
| C508c | 9, 13, 16, 17 |
| M289 | 8 |
| M291 | 9 |

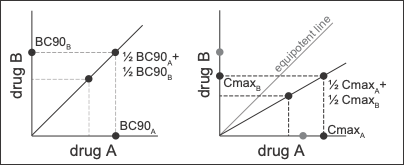

**Box S1. Explanatory schematic of equipotent and pharmacokinetics-informed dosing strategies.** For traditional equipotent dosing (left), a preliminary dose-centering experiment was performed to determine the IC or BC90 (concentration to achieve 90% inhibition or kill, depending on condition). Equipotent n-way combinations (where n = number of drugs in combination) were combined such that the seventh dose (out of 10) of the combination was a combination of the BC90s of the underlying singles divided by n. For pharmacokinetics-informed dosing (right), for each compartment (i.e., caseous or cellular) the *n*-way combinations were combined such that the highest dose was the Cmax of each drug in that compartment. Cmaxes were calculated from GranSim simulations (Methods).
